# An Integrative Approach for Fine-Mapping Chromatin Interactions

**DOI:** 10.1101/605576

**Authors:** Artur Jaroszewicz, Jason Ernst

## Abstract

Chromatin interactions play an important role in genome architecture and regulation. The Hi-C assay generates such interactions maps genome-wide, but at relatively low resolutions (e.g., 5-25kb), which is substantially larger than the resolution of transcription factor binding sites or open chromatin sites that are potential sources of such interactions. To predict the sources of Hi-C identified interactions at a high resolution (e.g., 100bp), we developed a computational method that integrates ChIP-seq data of transcription factors and histone marks and DNase-seq data. Our method, *χ*-SCNN, uses this data to first train a Siamese Convolutional Neural Network (SCNN) to discriminate between called Hi-C interactions and non-interactions. *χ*-SCNN then predicts the high-resolution source of each Hi-C interaction using a feature attribution method. We show these predictions recover original Hi-C peaks after extending them to be coarser. We also show *χ*-SCNN predictions enrich for evolutionarily conserved bases, eQTLs, and CTCF motifs, supporting their biological significance. *χ*-SCNN provides an approach for analyzing important aspects of genome architecture and regulation at a higher resolution than previously possible.

*χ*-SCNN software is available on GitHub (https://github.com/ernstlab/X-SCNN).

## 1. Introduction

Genome-wide maps of chromatin contacts are important for understanding genome architecture and gene regulation (Pope et al. 2014; Li et al. 2012; Fullwood et al. 2009). These contact maps also have implications to understanding the mechanism of disease-associated genetic variation (Lupiáñez et al. 2015; Won et al. 2016). Hi-C is an unbiased assay widely used for producing such genome-wide maps (Lieberman-Aiden et al. 2009). These maps are often represented with an *N*x*N* contact matrix, where *N* is the length of the genome divided by the chosen resolution. Within this matrix, subregions can be annotated as ‘peaks’ if the number of contacts within the subregion is significantly higher than expected (Rao et al. 2014; Ay et al. 2014; Pelossof et al. 2017). These peaks correspond to chromatin ‘loops’, where two loci are significantly closer to each other than regions in between them. Peaks enrich for promoters, enhancers, and cohesin-bound regions, which are often mediated by CTCF (Rao et al. 2014).

However, the resolution at which these peaks can be identified from Hi-C data is substantially larger than transcription factor (TF) binding or open chromatin sites, which can be considered potential sources of these interactions. The deepest human Hi-C sequencing experiment to date was performed on the GM12878 lymphoblastoid cell line with 3.6 billion reads generated (Rao et al. 2014) and led to a contact matrix at a 1kb resolution. However, interaction *peaks* were only reported at 5kb or 10kb resolution. Other cell types from the same study had peaks called at up to 25kb resolution, substantially larger than the 100-200bp resolution of TF binding and open chromatin sites. There are two major challenges with directly increasing resolution of Hi-C. Firstly, Hi-C is limited by the distribution of restriction sites (Naumova et al. 2012). Secondly, to increase resolution by a factor of *k*, one would need to increase the sequencing depth by *k*^2^.

We propose an alternative approach to obtain fine-resolution information in chromatin interaction peaks. Our approach is based on computationally integrating high-resolution ChIP-seq data of histone marks and TFs and DNase-seq data (Park 2009; Song & Crawford 2010). This is motivated by the observation that signal from such experiments shows specific patterns within interaction peaks such as pairs of CTCF sites or enhancer-promoter pairs (Rao et al. 2014). Our approach takes low resolution (e.g., 25kb) chromatin interaction peaks and uses DNase-seq and ChIP-seq data to predict at high resolution (e.g., 100bp) the source of each interaction. The approach is based on combining a Siamese Convolutional Neural Network (SCNN) trained to predict interactions with a feature attribution method to fine-map the interactions to their sources.

Limitations in the resolution of Hi-C have previously been recognized, and have inspired development of novel computational methods. For example, a transfer learning method was developed that learns from a high-resolution Hi-C map in one cell type to enhance the resolution of a Hi-C map in another cell type (Y. Zhang et al. 2018). While this was shown to effectively smooth noisy Hi-C data up to a 10kb resolution, it was not shown to be effective at finer resolutions. Other strategies have been proposed to enhance the resolution of contact maps genome-wide directly from Hi-C data (Carron et al. 2019; Cameron et al. 2018), but they are inherently limited to achieving at best restriction fragment length resolution. Other methods have been proposed that incorporate TF binding and epigenomic data to predict Hi-C data directly (Farré et al. 2018; S. Zhang et al. 2018). However, these methods are designed to make predictions at the resolution of the Hi-C data used for training, and not individual TF binding sites or open chromatin. By applying a feature attribution method, our approach makes predictions within these interacting regions, but at the finer resolution of DNase-seq and ChIP-seq data (∼100bp).

Other methods have aimed to solve related, but different, problems. Some methods have focused on using epigenetic data to predict specific aspects of chromatin structure genome-wide. For example, one method predicted the boundaries of Topologically Associated Domains (TADs) (Huang et al. 2015), and another reconstructed A/B compartments (Fortin & Hansen 2015). Other work aimed to predict promoter-enhancer interactions from epigenetic data, TF binding, or sequence data (Whalen et al. 2016; Cao et al. 2017; He et al. 2014; Roy et al. 2015; Singh et al. 2016), though the performance claims of some of these method in some cases has recently been challenged (Xi & Beer 2018). These methods differ from our proposed method in that their goal is to predict enhancer-promoter interactions, while we consider any type of Hi-C detected interaction, and our goal is to fine-map coarse, but detected, interactions.

In this paper, we first present our computational method, Chromatin Interaction Siamese Convolutional Neural Network (*χ*-SCNN, *χ* for the Greek letter Chi), to identify the likely sources of Hi-C identified interactions at high resolution. *χ*-SCNN leverages readily available high-resolution information in complementary data, specifically ChIP-seq and DNase-seq. We applied *χ*-SCNN to data from two cell types, and present a series of analyses quantitatively establishing the effectiveness of the approach. We also biologically characterize the fine-mapped positions. We expect *χ*-SCNN to be useful in the study of chromatin interactions.

## 2. Methods

Our method, *χ*-SCNN, uses a Siamese Convolutional Neural Network (SCNN) (Bromley et al. 1994; Koch et al. 2015) in conjunction with a feature attribution scoring method to fine-map called chromatin interactions. *χ*-SCNN first learns to discern called interactions from non-interactions using ChIP-seq data of histone marks and TF binding and DNase-seq data. It then performs fine-mapping by using Integrated Gradients (Sundararajan et al. 2017), a feature attribution method, to identify the pair of sub-loci that contribute most to the prediction of each interaction.

### Training Data

In *χ*-SCNN, each positive data point corresponds to an intra-chromosomal chromatin interaction peak. We applied *χ*-SCNN to peaks called from two Hi-C data sets: one from the lymphoblastoid cell line GM12878, and the other from the leukemia cell line K562 (Rao et al. 2014). We focused on these cell types because they also had DNase-seq and ChIP-seq data for many histone marks and TFs publicly available. For both of these cell types, we applied *χ*-SCNN to chromatin interaction peaks called by HiCCUPS at up to three different resolutions: 5kb, 10kb, and 25kb (Rao et al. 2014). For each peak, HiCCUPs chooses the finest resolution that surpasses a significance threshold. It called 9448 peaks in GM12878 and 6057 peaks in K562 at a False Discovery Rate (FDR) of 0.1.

Each negative data point corresponds to a sample from a distance matched, random genomic background. To form this background, *χ*-SCNN first computes the distribution of distances between interacting pairs observed in the positive training data, and then chooses random pairs of regions in the genome that match that distribution. We chose to use as many non-interacting pairs as observed interacting pairs to have a balanced dataset. We expected that when comparing interacting peaks to this negative background, *χ*-SCNN would learn which epigenetic and TF features differentiate peak regions from non-peak regions while controlling for distance-based effects.

### Feature Representation

Each interaction, whether a positive or negative data point, is represented by two matrices, one for each side of the interaction. The matrices are each of size *F* × *B*, where *F* is the number of features, and the number of bins *B = W/R*, where *W* is the width of the peak region, and *R* is the binning resolution. For each cell type, we used a set of previously uniformly processed DNase-seq and ChIP-seq data tracks from the ENCODE consortium^1^, where each track corresponds to one feature (ENCODE Consortium 2012). This resulted in 100 features for GM12878 and 148 features for K562. We later show that *χ*-SCNN is also effective with subsets of these features. We used *W* = 25kb for both GM12878 and K562 because this was the largest size peak called in these cell types, and it allowed for direct comparison of results (Rao et al. 2014). We use *R* = 100bp resolution for the binning resolution, yielding *B* = 25kb/100bp = 250 bins across each region.

Each ChIP-seq and DNase-seq track represents a normalized signal coverage (Hoffman et al. 2013). For each track, we first averaged the values within each bin. We then added 1 to each value and then performed a log_2_-transformation to make the training more robust to extreme outliers. Finally, because only the relative orientation of the regions is relevant and not the specific strand they are on, we took each pair of matrices and reversed their direction to go from 3’ to 5’ in addi tion to 5’ to 3’, and then added them to the dataset. This was a simple way to increase the size of our training set and reduce overfitting.

### Neural Network Architecture

*χ*-SCNN uses a Siamese Convolutional Neural Network (SCNN, **Fig. 1**) (Bromley et al. 1994; Koch et al. 2015). SCNNs are composed of two identical subnetworks with shared parameters, which are later joined to make a prediction. They are often used to determine if two objects are similar, but in this application, it determines whether two genomic regions interact based on their epigenetic and TF signatures, which may be dissimilar. Each subnetwork processes its input separately, after which the dense (fullyconnected) layer integrates data from these subnetworks. Each subnetwork is composed of a compressing encoding layer, followed by a convolutional layer, then a global max-pooling layer. Data from the global max-pooling layers is then passed to a dense layer and, finally, a logistic regression layer, which calculates a probability of the two regions interacting. *χ*-SCNN uses a ReLu nonlinear transformation, defined as ReLu(*x*) = max(0, *x*), after the encoding, convolution, and dense layers.

**Figure 1.**
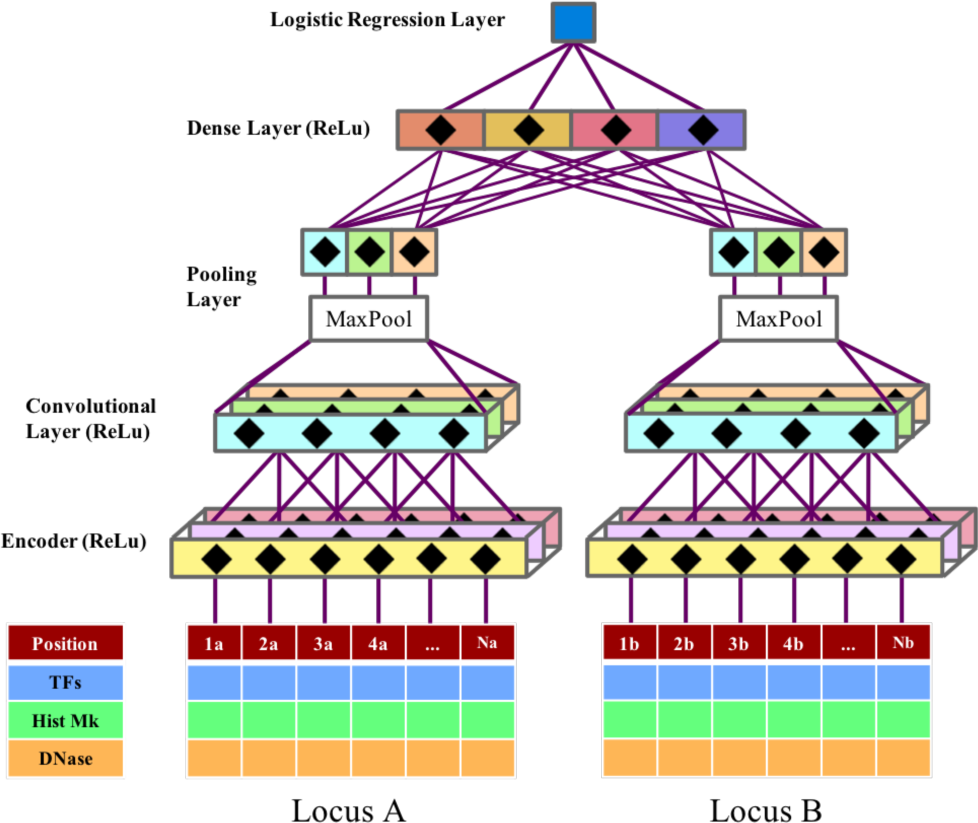
The structure of the SCNN in *χ*-SCNN. Two data matrices are passed in parallel through an encoding layer, convolutional layer, global max pooling layer, a dense layer, and finally, a logistic regression layer. The encoder, convolutional, and dense layers use a ReLu activation function. The two subnetworks are identical (all weights are shared) until the dense layer.

### Encoder

The encoder projects a high-dimensional space (*F*x*B*) to a lower one (*K*_*Enc*_x*B*), where *K*_*Enc*_*< F* is the number of encoder kernels. Encoders make neural networks easier to optimize and more difficult to overfit, and have a similar objective to other dimensionality reduction approaches such as Principal Component Analysis (PCA) (Bourlard & Kamp 1988; Chicco et al. 2014), except that they are further tuned after initialization. Here, *χ*-SCNN initializes the encoder by pre-training an autoencoder (an encoder-decoder pair) (Ballard 1987) on interacting regions in the training data, then transfers the learned encoder weights to the final SCNN. The autoencoder has a width of 1 bin, meaning that it is applied to each position independently, and only has one hidden layer to keep the number of parameters low and prevent overfitting.

### Convolutional layer

Following the encoder, the convolutional layer slides a matrix of *K*_*Enc*_*xC* values across the entire matrix of size *K*_*Enc*_*xB* for each of the *K*_*Conv*_ kernels. At each of the *D = B*-*C+*1 sub-matrices of size *K*_*Enc*_*xC* in the region, it calculates the element-wise product of the data with each of *K*_*Conv*_ kernels, followed by summation. This produces a matrix of *K*_*Conv*_*xD* values. The computed values are then passed through a ReLu transformation, which sets negative values to zero. Intuitively, the convolution layer is used to find local spatial patterns of signal in the ChIP/DNase-seq data, such as co-binding of several TFs or a promoter followed by an actively transcribed region. Importantly, it does not make any assumptions about the specific positions of patterns in a region, a useful characteristic for our application, as the interaction source is expected to be in different positions for different interactions.

### Global max-pooling layer

The global max-pooling layer takes as input the *K*_*Conv*_*xD* matrix output by the convolutional layer, and then outputs the maximum value for each of the *K*_*Conv*_ rows.

### Dense layer

The outputs of the global max-pooling layers from two subnetworks are then integrated at the dense layer. Each of the *K*_*Dense*_ kernels in the dense layer has access to every value from the global max-pooling layer computed across each of the subnetworks. It multiplies these values by learned weights, adds a bias term, and outputs a vector of size *K*_*Dense*_. This layer finds which signal profiles are compatible with each other. For example, it can potentially learn that regions with enhancer-like elements tend to co-occur with promoter-like elements.

### Logistic regression layer

The final layer is the logistic regression layer, which takes the *K*_*Dense*_ values output from the dense layer, multiplies them by learned weights, and passes their sum with a learned bias term through the logistic function. The logistic layer returns a probability between 0 and 1, corresponding to the model’s confidence that a sample is positive.

### Training and hyperparameter search

We implemented *χ*-SCNN using Keras, a Python neural network library built on top of TensorFlow (Abadi et al. 2016). The autoencoder is pretrained to optimize a mean squared logarithmic error loss function, which is appropriate for continuous data. For binary classification, the whole SCNN uses a binary cross-entropy loss function. Both use Stochastic Gradient Descent using the ADADELTA optimizer (Zeiler 2012). Chromosomes 8 and 9 are withheld for validation after each epoch of training, and chromosome 1 for final testing evaluation. We performed a random search to select a combination of hyperparameters (Bergstra & Bengio 2012). Specifically, we searched for the width of the convolutional filter (*C*), the number of kernels for the autoencoder (*K*_*Enc*_), convolutional layer (*K*_*Conv*_), and dense layer (*K*_*Dense*_), the type and strength of regularization for all trained parameters, and the dropout magnitude (Srivastava et al. 2014) (**Supp. Table 1**). For each dataset, we tried 60 random combinations of hyperparameters, as it yields at least a 95% probability of the performance being within the top 5% of hyperparameter choices^2^. We chose the hyperparameter combination that achieved the best AUROC on validation data, and we report the test data AUROC and AUPRC for chromosome 1 (**Supp. Table 1**). We note that different applications of *χ*-SCNN can lead to the selection of different hyper-parameter combinations.

**Table 1.** Enrichments of functional elements as compared to the genome. *χ*-SCNN is compared to baseline methods: (1) logistic regression, (2) randomly guessing a position within the peak region, and (3) using the 1kb position with the highest signal (available only in GM12878). Values given in parentheses are relative to randomly guessing within the called interaction peak.

| Cell type | GM12878 |  |  |  | K562 |  |  |
| --- | --- | --- | --- | --- | --- | --- | --- |
| | χ-SCNN | Logistic Regression | Random in peak | Hi-C 1kb Signal | χ-SCNN | Logistic Regression | Random in peak |
| GERP++ elements | **5.82**<br>(2.45) | 5.17<br>(2.26) | 2.38<br>(1.00) | 2.29<br>(0.96) | **5.05**<br>(2.21) | 4.17<br>(1.83) | 2.28<br>(1.00) |
| eQTL (matched) | **5.63**<br>(1.62) | 4.15<br>(1.20) | 3.47<br>(1.00) | 3.18<br>(0.92) | **4.34**<br>(1.05) | 4.08<br>(0.99) | 4.14<br>(1.00) |
| CTCF motifs | **768.53**<br>(116.09) | 676.78<br>(102.23) | 6.62<br>(1.00) | 6.44<br>(0.97) | **725.58**<br>(102.48) | 583.32<br>(82.39) | 7.08<br>(1.00) |

Finally, after selecting the optimal hyperparameter combination, *χ*-SCNN is retrained using all peaks except those on chromosomes 8 and 9, which are used as a stopping condition for training. This model is used for fine-mapping and all subsequent analyses.

### Fine-Mapping

After training the SCNN, *χ*-SCNN fine-maps peaks using a feature attribution method to score each position within the peak region. Feature attribution methods were developed to help explain why a machine learning model made a specific prediction. For example, in the context of image recognition, it can be used to determine which pixels of an image contributed the most to the image’s classification; if an image is predicted to contain a cat, it would be expected to highly score areas around the whiskers and ears, but ignore irrelevant background.

*χ*-SCNN uses a feature attribution method called Integrated Gradients (Sundararajan et al. 2017) to apply the same methodology to its predictions, wherein ‘pixels’ correspond to bins in the input matrices. We chose to use Integrated Gradients because of its simplicity in assumptions and implementation. As in the original application of Integrated Gradients, to determine the importance of a pixel with multiple individual features (RGB values), *χ*-SCNN determines the total importance for a bin by summing the importances of all features at that bin (**Fig. 2**).

**Figure 2.**
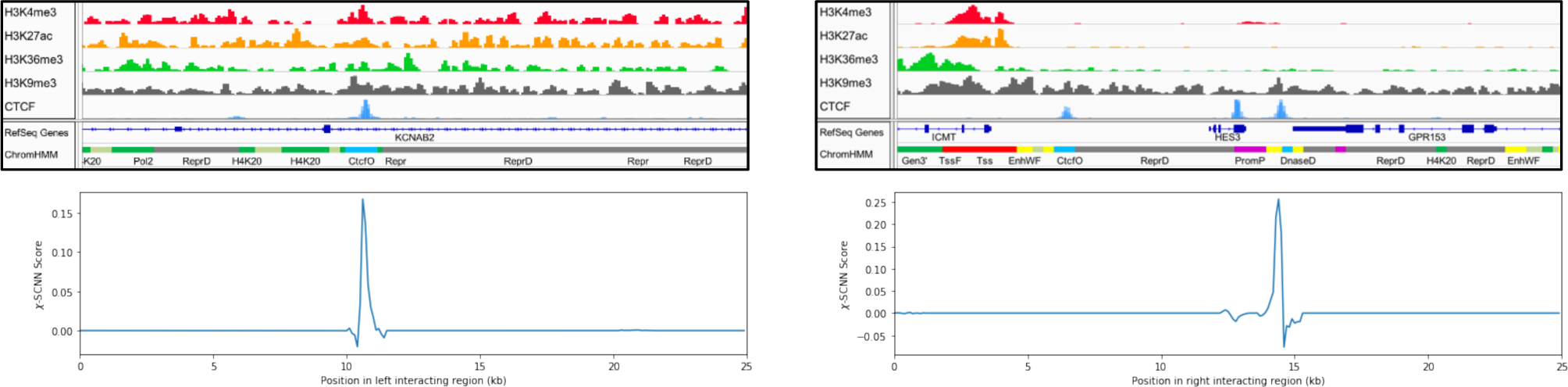
An example of a fine-mapped peak. The left and right sides correspond to the two sides of the interactions. The top images show tracks for H3K4me3, H3K27ac, H3K36me3, H3K9me3, and CTCF. The bottom images show *χ*-SCNN’s fine-mapping score for each position in the region. There is a sharp peak on the left corresponding to a CTCF peak, and in the right region, *χ*-SCNN is given three CTCF peaks, and predicts one of them.

The feature importance scores that Integrated Gradients assigns are roughly equal to the output probability difference when setting that feature to a baseline of 0 (which corresponds to no signal) (Sundararajan et al. 2017). A score of *s* > 0 at some bin means that setting the data in that bin to 0 would decrease the calculated probability of the two regions interacting by approximately *s*. Conversely, if *s* is negative, setting the data in the bin to 0 would increase the probability of interaction by *s*. For each side of an interaction, we took the position with the highest overall score as the ‘fine-mapped’ peak, and used these positions for all subsequent analyses and validation.

## 3 Results

### 3.1 *χ*-SCNN is highly predictive of interactions

Before fine-mapping called interactions, we first established that the SCNN of *χ*-SCNN is effective at discriminating between positive and negative interactions. We note that this is a necessary, though not sufficient, condition for fine-mapping called interactions. We conducted the evaluations on a withheld test set of interactions on chromosome 1, which was not used for training the SCNN or selecting hyper-parameters. The SCNN achieved a high Area Under the Receiver Operator Characteristic (AUROC) curve for predicting interactions in GM12878 and K562, 0.96 and 0.98, respectively. We also evaluated the area under the Precision-Recall curves (AUPRC), and obtained values of 0.96 and 0.97 for GM12878 and K562, respectively (**Supp. Table 1**). We note that the AUPRC depends on the ratio between positive and negative samples, and since we are considering balanced data it is expected to be higher here than if predicting genome-wide. We emphasize, however, that our goal is not to predict interactions genome-wide, but rather to fine-map called interactions.

### 3.2 *χ*-SCNN fine-mapping predictions are reproducible

Having established that *χ*-SCNN could effectively discriminate positive from negative interactions, we next sought to establish that *χ*-SCNN’s fine-mapping method was reproducible. For each cell type, we took the corresponding set of HiCCUPs peaks calls and split by chromosome into two non-overlapping sets of approximately equal size; one set was composed of interactions on odd chromosomes, and the other on even chromosomes and chromosome X. We did not create a third withheld set because the typical application of *χ*-SCNN is training and fine-mapping on the same set of peaks. We trained two separate models, one on each split set. We then fine-mapped all the interactions and calculated the fine-mapping concordance by calculating Euclidean distance on a 2D grid between the fine-mapped peaks (**Supp. Fig. 1**). We found that 90% and 87% of interactions fine-mapped within 100bp in any direction for GM12878 and K562, respectively, as compared to an expected 0.01% by chance, and this concordance further increased at more relaxed distance thresholds. We note that each of the datasets in this analysis was roughly half the size of the full dataset, and thus the results should be considered a lower bound of expected reproducibility.

### 3.3 *χ*-SCNN fine-mapped predictions recover original Hi-C peaks after extension

Having established *χ*-SCNN fine-mapping predictions are reproducible, we next sought evidence that they are also accurate. As we do not have Hi-C interaction peak calls available at the resolution of *χ*-SCNN predictions, we instead evaluated how well *χ*-SCNN predictions can identify the original called peaks when provided those peaks after extending their boundaries.

For each peak narrower than 25kb (i.e., 5kb or 10kb), we extended the boundaries of each side of an interaction uniformly in both directions to produce a 25kb peak; for example, if one side of a 5kb interaction involved the interval 140kb-145kb, we extended it to the interval 130kb-155kb. Together with peaks originally called at 25kb, we extracted ChIP-seq and DNase-seq data from these 25kb regions and applied *χ*-SCNN. We then evaluated how often the fine-mapping fell in the center 5kb region.

We found that for 5kb HiCCUPs peaks in K562 extended to 25kb, *χ*-SCNN fine-mapping predictions were, as expected, frequently found in the center 5kb region (33% of peaks, 8.2 fold enrichment compared to random guessing, p-value < 0.001, binomial test) (**Fig. 3a**). Similarly, fine-mapping predictions of 10kb HiCCUPS peaks in K562 extended to 25kb had a strong enrichment in the center 5kb (4.3 fold enrichment, p-value < 0.001) (**Fig. 3b**). Peaks that were originally called at 25kb had a much smaller enrichment in the center cell (1.5 fold enrichment, p-value < 0.001) (**Fig. 3c**). This smaller enrichment was expected, since the true peak source is more likely to fall anywhere within the 25kb region than for peaks called at a finer resolution. We also applied the same evaluations to GM12878 and found that it performed slightly better with 8.9 and 4.5 fold enrichment for 5kb and 10kb peaks, respectively. These results show that *χ*-SCNN strongly enriches for recovering finer resolution peaks after extending the peaks to be 25kb (**Supp. Table 2**, **Supp. Fig. 2**).

**Figure 3.**
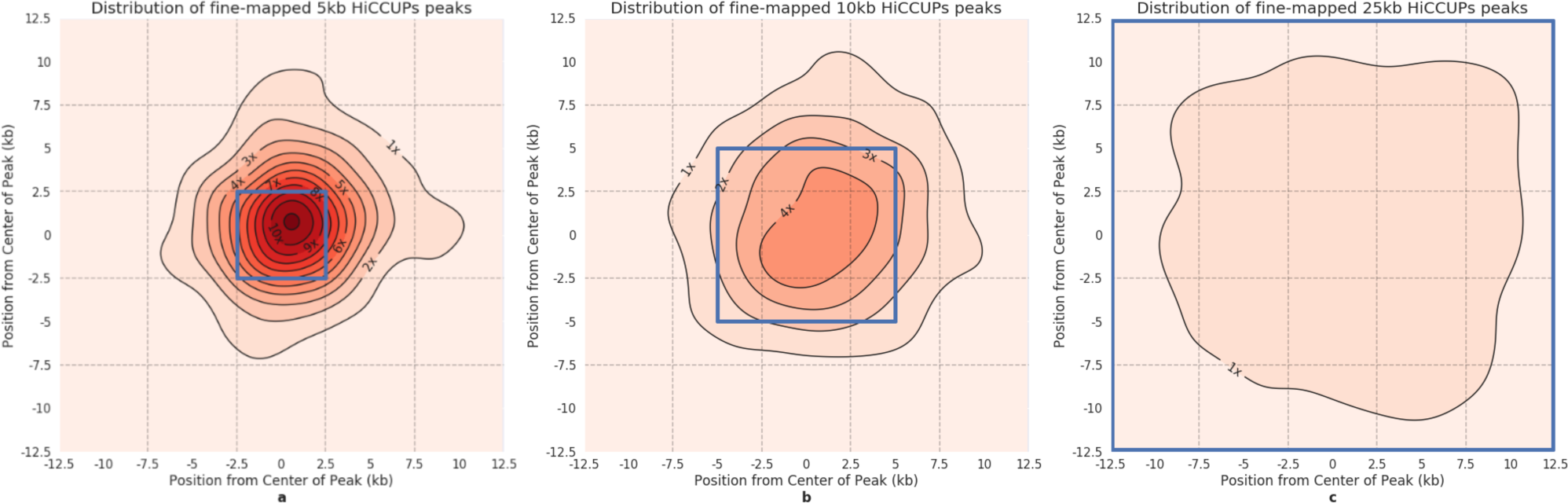
Distribution of fine-mapping predictions for different size HiCCUPs peaks. Kernel Density Estimation (KDE) plots showing the distribution of *χ*-SCNN’s fine-mapping predictions within K562 peaks after extending the original peak equally in both directions to form a 25kb peak. To generate plots, we used the ‘jointplot’ function with the KDE option in Python’s Seaborn package. **(a)** For 5kb interaction peaks extended to 25kb, fine-mapped positions are strongly concentrated around the original 5kb peak (center blue bin). Enrichment in center 5kb bin is 8.2 fold compared to random guessing. **(b)** For 10kb peaks extended to 25kb, fine-mapped positions are concentrated in the original 10kb peak (center blue bin). Enrichment in center 5kb bin is 4.2 fold. **(c)** Fine-mapped positions are not concentrated in any specific region in interactions called at 25kb. Enrichment in center 5kb bin is 1.5 fold. The positive direction on the axes points toward the exterior of the interactions. The mode of the 5kb peak plot is shifted toward the positive direction, meaning that fine-mapped peaks are most likely to be approximately 1kb further out than the center of the called originally called peak. Similar plots for GM12878 can be found in **Supp. Fig. 2**.

We also applied *χ*-SCNN to 5kb peaks after extending unevenly, specifically extending the original 5kb peak by 20kb on one side and holding the other side fixed. We found that *χ*-SCNN performed similarly in peak recovery as when extending uniformly (8.1 vs. 8.2 fold enrichment in K562 and 8.9 fold for both in GM12878).

When extending evenly, a large percentage of *χ*-SCNN fine-mapping predictions for extended 5kb peaks did not map to the center bin, but to one of the four directly adjacent bins (39%, 2.4 fold enrichment, p-value < 0.001 for both cell types). Many of these off-center predictions could be expected to be the true source, as the original HiCCUPS predictions are based on noisy Hi-C data, which can lack the resolution to differentiate between interaction sources near the boundary of two 5kb cells.

Interestingly, we found that the mode of *χ*-SCNN fine-mapped peaks was not at the center of the original 5kb peak, but about 1kb toward the exterior of the interaction on each side (**Fig. 3a, Supp. Fig. 2a**). Conversely, the HiCCUPs peak calls were likely to be called the opposite way, approximately 1kb inward from the predicted source of the interaction. This observation suggests a bias in either the Hi-C experiment or the HiCCUPs peak caller toward the interior of a looping interaction as compared to the fine-mapped peak, as *χ*-SCNN cannot learn a spatial bias or preference except for possible boundary effects. The difference could be explained by the strong effect of distance on Hi-C signal, as pairs of regions that are closer to each other tend to exhibit higher signal.

### 3.4 *χ*-SCNN better recovers original Hi-C peak after extension than baseline approaches

We compared the performance of *χ*-SCNN at recovering the original 5kb and 10kb Hi-C peaks to several baselines (**Supp. Table 2**). The first set of baselines consisted of considering each DNase-seq and ChIP-seq track separately and fine-mapping to the position with the highest signal. This included 100 tracks for GM12878 and 148 tracks for K562. In cases where there were multiple features corresponding to the same histone mark, TF, or DNase-seq, we also created a baseline prediction by averaging the signals across those features, and we used this for the reported enrichments. We note that the best performing of these baselines only provides an upper bound on expected fine-mapping performance when selecting a single track or feature average, as in practice there is no guarantee the selection made would be optimal for fine-mapping.

We found that several TFs, notably CTCF and the cohesin marks RAD21 and SMC3, had high enrichment for recovering the original 5kb and 10kb HiCCUPs peaks (**Supp. Table 2**), consistent with their previously reported high enrichment in interactions (Rao et al. 2014), but were all less than *χ*-SCNN’s predictions. Combining counts from both 5kb peaks and 10kb peaks, *χ*-SCNN outperformed all other tracks (p-value < 0.05, two-proportions z-test). All other ChIP-seq tracks and DNase had lower performances, at most 6.4 and 5.2 in GM12878 and K562, respectively, as compared to the 8.9 and 8.2 fold enrichment for *χ*-SCNN (**Fig. 4**).

**Figure 4.**
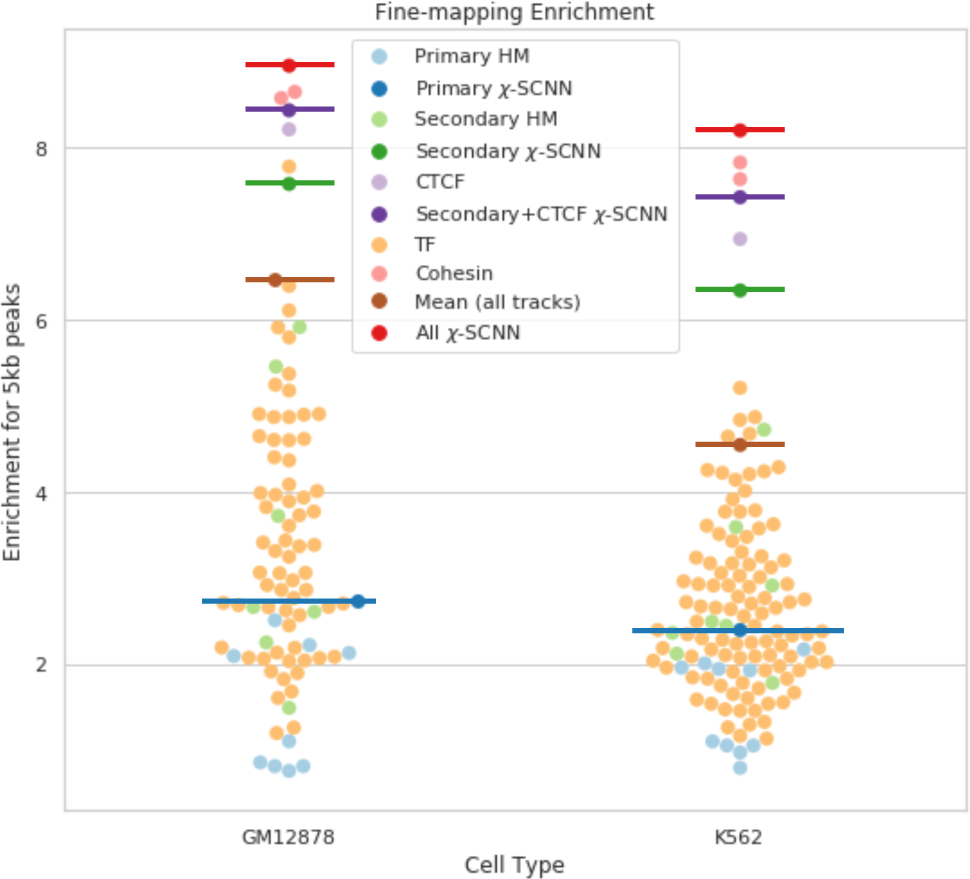
5kb peak fine-mapping performance for *χ*-SCNN and baseline methods. Fine-mapping performance using individual features is marked with points, and methods integrating multiple features are emphasized with horizontal bars. Light blue points and the dark blue bar correspond to ‘primary’ histone marks and *χ*-SCNN trained on these marks, respectively. Similarly, light green points and the dark green bar correspond to ‘secondary’ marks. CTCF, in lavender, performs well, but *χ*-SCNN trained on ‘secondary’ marks and CTCF together performs better. Cohesion sub-units RAD21 and SMC3, in pink, are the best performing single marks; however, *χ*-SCNN trained on all available features, in red, shows greater enrichment than any individual mark. All other TFs, in orange, perform similarly to histone marks. Finally, a baseline method of averaging all features is marked with a brown bar.

Another baseline we evaluated was predicting based on averaging all ChIP-seq and DNase signal tracks and then taking the position with highest average signal. For the 5kb evaluation, this had a fold enrichment of 6.5 and 4.5 in GM12878 and K562, respectively, which was significantly less than *χ*-SCNN’s enrichments (p-value < 0.001) (**Fig. 4**).

Finally, we compared our fine-mapping predictions to a logistic regression model trained to distinguish between true interactions and random ones. To train in such a way as to fine-map to 100bp resolution, we randomly sampled 100,000 pairs of 100bp positions from the positive interactions. We used the same methodology for the negative set, except sampling from randomly generated, distance controlled genomic pairs. We trained the logistic regression model with default parameters using the Python package scikit-learn. To fine-map, we took each pair of 100bp regions for each peak and returned the pair that yielded the highest probability. For the 5kb evaluation, the logistic regression model achieved 8.2 fold and 7.0 fold enrichments for GM12878 and K562, respectively. These were also significantly lower than *χ*-SCNN’s enrichments (p-value < 0.001, two-proportions z-test). *χ*-SCNN was also faster in fine-mapping, as it is linear in complexity with regard to the number of bins, while logistic regression is quadratic in complexity, as each pair of 100bp bins must be explicitly evaluated.

### 3.5 *χ*-SCNN outperforms baseline approaches in recovering relevant external annotations

We next analyzed the enrichment of *χ*-SCNN’s fine-mapping predictions and baseline approaches for several external annotations. The external annotations considered are defined at or near base pair resolution and are suggestive of functionally relevant positions. Specifically, we considered: (1) Evolutionarily conserved bases, as this is a relatively unbiased annotation of likely functionally relevant positions. We used GERP++ elements to define these (Davydov et al. 2010); (2) expression Quantitative Trait Loci (eQTL) variants, as they provide evidence a position may affect expression of genes at distal loci, and transcriptional regulation has been shown to be associated with chromatin contacts (Li et al. 2012). The eQTL annotations were obtained from GTEx (The GTEx Consortium 2017). We used EBV-transformed lymphocytes and whole blood, as these cell types are closely related to GM12878 and K562, respectively; (3) CTCF motifs annotations (Kheradpour & Kellis 2014), as their importance in loop interactions has previously been established (Rao et al. 2014; Sanborn et al. 2015). We expected that more accurate fine-mapping predictions would show overall greater enrichment for these annotations.

For each of these annotations, we calculated the average overlap of bases between the annotation and *χ*-SCNN’s 100bp fine-mapped predictions. We then calculated the enrichment of these overlaps relative to the entire genome. We compared these enrichments to enrichments from (1) randomly guessing within peak regions, (2) predictions from the logistic regression baseline, and (3) directly using the GM12878 1-kb Hi-C data signal, the finest resolution Hi-C data available for humans (Rao et al. 2014). We performed this final comparison only in GM12878, as 1kb resolution data is not available for K562.

To make a fine-mapping prediction directly from Hi-C signal, we first took the number of reads in each 1kb by 1kb Hi-C contact matrix cell for the corresponding peak. We then divided by the KR normalization vectors as in (Knight & Ruiz 2012; Rao et al. 2014), and took the cell with the highest normalized signal. We found that *χ*-SCNN outperformed the baseline methods in all comparisons (**Table 1**). Surprisingly, directly using the 1-kb Hi-C data did not provide any additional predictive power in recovering GERP++ elements, eQTLs, or CTCF motifs over randomly guessing within the peak region. This suggests that 1kb Hi-C signal does not have additional information for their recovery beyond the 5-25kb interaction peak, and highlights the value of integrating epigenomic or TF binding data to make finer resolution predictions.

### 3.6 Fine-mapped positions show distinct chromatin state enrichments

To gain insight into the type of locations that are predicted to be the source of interactions, we analyzed *χ*-SCNN’s predictions relative to a 25-state ChromHMM model from the ENCODE integrative analysis (Ernst & Kellis 2012; Ernst & Kellis 2013; Hoffman et al. 2013). For each H iCCUPs peak in a given cell type, we took the highest-scoring 100bp sub-region on each side and found the corresponding pair of ChromHMM annotations. We counted the number of fine-mapped sites found for each ChromHMM state, then added a pseudocount of 1 to the count of each state. We normalized this to find a frequency of each state and compared it to average genomic frequency to compute the fold enrichment, and then took the log_2_ of this value (**Fig. 5, Supp. Fig. 3**). The most enriched state was ‘CtcfO’, a state associated with CTCF binding in open chromatin regions, with a fold enrichment of 267 and 224 in GM12878 and K562, respectively.

**Figure 5.**
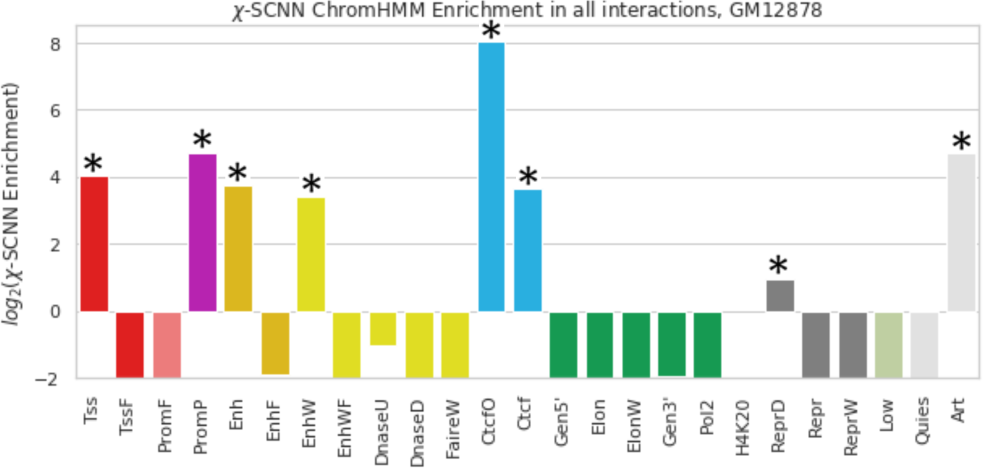
Enrichment of ChromHMM states of regions predicted by *χ*-SCNN. Shown is the log_2_ fold enrichments of ChromHMM state for *χ*-SCNN fine-mapped positions in GM12878 interactions. Any log_2_ fold depletions less than −2 (1/4 fold) were truncated. Significant enrichments (adjusted p-value < 0.001, binomial test) are marked by an asterisk. Across all interactions, the ChromHMM state ‘CtcfO’ is most enriched, at 267 fold genome-wide enrichment. Analogous enrichment plots separately for CTCF-associated and non-CTCF-associated interactions can be found in **Supp. Fig. 3** as well as for K562.

Besides the ‘CtcfO’ state, we also found notable enrichment for states associated with transcription start sites (‘Tss’, 17 and 15 fold enrichment for GM12878 and K562), poised promoters (‘PromP’, 26 and 35 fold enrichment), enhancers (‘Enh’, 14 and 12 fold enrichment), weak enhancers (‘EnhW’, 11 and 14 fold enrichment), CTCF binding without open chromatin (‘Ctcf’, 13 and 10 fold enrichment), and the artifact state (‘Art’, 27 and 24 fold enrichment).

### 3.7 *χ*-SCNN is effective using a limited set of features

In applying *χ*-SCNN to GM12878 and K562, we used more input features than are typically available in most cell types. To estimate the expected performance of *χ*-SCNN for cell types with more limited data, we evaluated its performance using subsets of features. Our basic feature set is composed of ChIP-seq data for ‘primary’ histone marks: H3K27me3, H3K36me3, H3K4me1, H3K4me3, H3K9me3, and H3K27ac, which are available for 98 cell and tissue types from Roadmap Epigenomics. We then extended this ‘primary’ feature set with a ‘secondary’ set by adding the histone marks H3K4me2, H3K9ac, H4K20me1, H3K79me2, and H2A.Z, in addition to DNase-seq. We chose this feature set as these features were all were deeply mapped by either the ENCODE or Roadmap Epigenomics projects, and they are available as imputed data for 127 cell and tissue types (Ernst & Kellis 2015). As CTCF is also available for many cell types, we also tried adding CTCF to the secondary set. Finally, we compared results from these three sets to results achieved by using all data.

We first evaluated the performance of the SCNN at discriminating between positive and negative interactions using subsets of features. We found that when using only primary marks, the performance was reasonably high (AUROCs of 0.80 and 0.83 for GM12878 and K562, respectively). The performance increased substantially by adding the secondary set of features, (AUROCs of 0.94 and 0.95) and a smaller improvement when further adding CTCF (AUROCs of 0.96 and 0.97), close to the performance using all the marks (AUROCs of 0.96 and 0.98).

We then evaluated the performance in peak recovery after extending 5kb HiCCUPs peaks to 25kb using the same subsets of marks. Using only primary marks yielded a fold enrichment in the center 5kb window of 2.7 and 2.4 for GM12878 and K562; adding the secondary set had a larger enrichment of 7.6 and 6.3; adding

CTCF to this set yielded 8.4 and 7.4 enrichment, whereas using all marks had the largest enrichment at 8.9 and 8.2 (**Fig. 4**).

### 3.8 *χ*-SCNN fine-mapping reveals CTCF-associated and non-CTCF-associated interactions

Because CTCF is considered a crucial component in forming and maintaining chromatin interactions (Sanborn et al. 2015), we investigated how *χ*-SCNN predictions relate to CTCF binding. To investigate this we first downloaded peak calls for ChIP-seq data of CTCF^3^ and took the intersection of peak calls from different laboratories following the procedure of (Rao et al. 2014). We compared CTCF peaks with HiCCUPs peaks, and found that most interacting regions overlapped at least one CTCF peak (72.0% for GM12878, and 83.3% for K562), largely consistent with previous findings (Rao et al. 2014). This is lower than the 86% and 88.1% previously reported, which was found by extending peaks to 15kb before finding CTCF peak overlap (Rao et al. 2014). When extending HiCCUPs peaks to 25kb as in this application, the number rises to 95.3% and 95.2% for GM12878 and K562, respectively.

We observed that 34% and 44% of regions involved in an interaction contained multiple CTCF peaks in GM12878 in K562, respectively. We investigated whether *χ*-SCNN can better identify original 5kb interacting peaks after extending the peak to 25kb, relative to two baselines: (1) choosing the CTCF peak with the highest signal, and (2) choosing a CTCF peak at random. We evaluated fine-mapping on a one-dimensional axis instead of a two-dimensional grid as in (**Fig. 3**), as we were only evaluating individual sides of interactions that had multiple CTCF peaks. We found that using *χ*-SCNN with all marks had a 2.8 and 2.7 fold enrichment in recovering the original 5kb peaks for GM12878 and K562, respectively. This was significantly greater than the enrichments from choosing the peak with the highest CTCF signal, 2.5 and 2.3, and from randomly guessing a CTCF peak, 2.0 enrichment for both cell types (p-value < 0.001, two-proportions z-test).

We then separated all of *χ*-SCNN fine-mapping predictions into two sets: ‘CTCF-associated’, those that overlapped the union of all CTCF peaks (92.5% and 93.8% for GM12878 and K562, respectively), and the remaining ‘non-CTCF-associated’ that did not overlap any CTCF peaks. We chose to look at the union of CTCF peaks to get a more confident set of peaks that did not overlap CTCF. Of the non-CTCF-associated interactions, 37.2% in GM12878 and 22.2% in K562 had a CTCF peak not at the fine-mapped position, but elsewhere in the broader interacting region, meaning that *χ*-SCNN does not simply predict based on CTCF.

Finally, we compared chromatin state enrichments between CTCF-associated and non-CTCF-associated interactions. We found that CTCF-associated interactions were mostly frequently mapped and highest enriched in the ‘CtcfO’ state (280 fold enrichment for GM12878 and 236 fold enrichment for K562). In GM12878, non-CTCF-associated interactions were most likely to map to the ‘Tss’ and ‘Enh’ states (52 and 47 fold) (**Fig. 5**). In K562, they showed large enrichments for the ‘Tss’, ‘Enh’, and ‘Ctcf’ states (29, 14, and 12 fold) (**Supp. Fig. 3**). We also saw substantial enrichments for the ‘FaireW’ and ‘Art’ states, but these accounted for only 13.8% of interactions compared to 24.6% for the ‘Tss’ and ‘Enh’ states combined. These results suggest a substantial contribution from promoters and enhancers among the non-CTCF-associated interactions.

### 3.9 Limited interaction specificity found with shuffled background

The distance-matched, random genomic background allows *χ*-SCNN to learn signatures of locations involved in interactions in general, but not necessarily *pairwise* signatures for pairs of interacting loci. We investigated whether epigenetic and TF data could inform which *pairs* of regions interact given all interacting regions. To do this, we modified the negative training dataset to control for per-locus signal of all input tracks. Specifically, we generated a negative training dataset where instead of randomly sampling two random genomic loci, we shuffled interactions. In other words, we took each interacting locus and paired it with a different one at random; each region that was part of an interaction was now paired with a region from a different interaction. We followed the same procedure for hyperparameter search, training, and testing as with the genomic background. We found that models trained on Hi-C peaks in GM12878 and K562 achieved AUROCs of 0.64 and 0.70, respectively. This suggests there is some detectable pairwise epigenetic and TF binding signal predictive of interactions, but because of the relatively low separability of true and shuffled interactions, we were unable to robustly characterize this pairwise signal.

## 4. Discussion

We developed *χ*-SCNN, a method for fine-mapping coarse Hi-C interactions to their sources by leveraging high resolution ChIP-seq and DNase-seq data. The method applies an SCNN to learn epigenomic signatures of pairs of interactions. We then analyzed each pair of interactions using a feature attribution method, Integrated Gradients, to identify the positions that are most informative to the prediction of the interaction, and thus can be inferred to be the ‘fine-mapped’ peak.

We applied *χ*-SCNN to data from two cell types and demonstrated that it effectively identifies original Hi-C peaks after extending them. We demonstrated that *χ*-SCNN has higher enrichment than using the signal of any single mark alone or the average of all them. We showed that *χ*-SCNN predictions have greater enrichment for evolutionarily conserved bases, eQTLs, and CTCF motifs than several baseline comparisons, which suggests greater functional relevance of *χ*-SCNN predictions. The fine-mapped loci also strongly enrich for primarily CTCF-associated chromatin states, which is expected based on existing knowledge (Rao et al. 2014; Sanborn et al. 2015), and also highlighted enhancer and promoter states associated with non-CTCF-associated interactions.

We note that using our framework, we can apply alternative background models to potentially detect subtler, but still potentially biologically relevant signal. Specifically, we investigated an alternative ‘shuffled’ background, which was a way to identify if there was pairwise epigenetic signal that differentiated interactions from each other, as opposed to identifying signals associated with interactions in general. However, when training against this shuffled background, we saw limited predictive power, suggesting limited pairwise signal, consistent with previous observations in the context of predicting enhancer-promoter interactions (Xi & Beer 2018).

We demonstrated that *χ*-SCNN is effective using only a subset of features that is available for many cell types (ENCODE Consortium 2012; Roadmap Epigenomics Consortium et al. 2015). While we focused on Hi-C here, an avenue for future investigation would be to apply and evaluate *χ*-SCNN on other types of interaction data such as Promoter-Capture Hi-C, ChIA-PET, or Hi-ChIP data (Mifsud et al. 2015; Fullwood & Ruan 2009; Mumbach et al. 2016). We expect *χ*-SCNN predictions to serve as a resource to better understand chromatin interactions and non-coding variants relevant to disease.

## Acknowledgements

We would like to acknowledge Anshul Kundaje, Sriram Sankararaman, and members of Jason Ernst’s lab for their valuable input during the method development process.

## Footnotes

1 Data available at http://ftp.ebi.ac.uk/pub/databases/ensembl/encode/integration_data_jan2011/byDataType/signal/jan2011/bigwig/

2 The probability of a model trained with a random combination of hyperparameters not being in the top 5% of all combinations is 0.95. For *n* combinations of hyperparameters, the probability of none of them being in the top 5% of all combinations is (0.95)^*n*^, which is less than 0.05 for n≥59.

3 Data available at http://ftp.ebi.ac.uk/pub/databases/ensembl/encode/integration_data_jan2011/byDataType/peaks/jan2011/spp/optimal/

