## supplementary figures and tables for "An Integrative Approach for Fine-Mapping Chromatin Interactions"

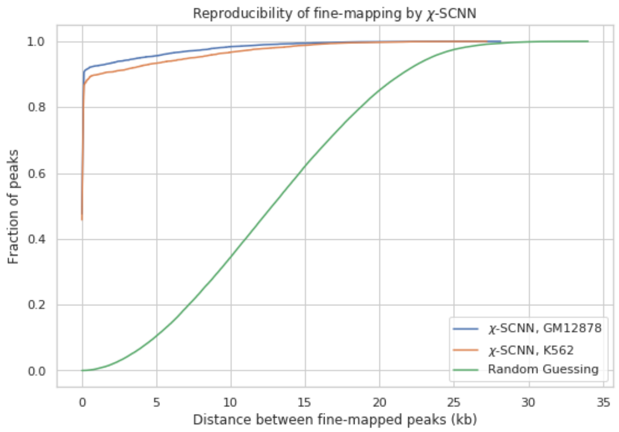

**Supplementary Figure 1. Reproducibility of fine-mapped peaks.** The plot shows as a function of Euclidean distance (x-axis) the fraction of fine mapped that fell within that distance (y-axis) from two different χ-SCNN models for both GM12878 and K562. 90% and 87% of fine-mappings fall within 100bp in either direction for GM12878 and K562, and 93% and 90% of peaks fall within 1kb. Also shown is a baseline of random guessing that is shared for the two cell types.

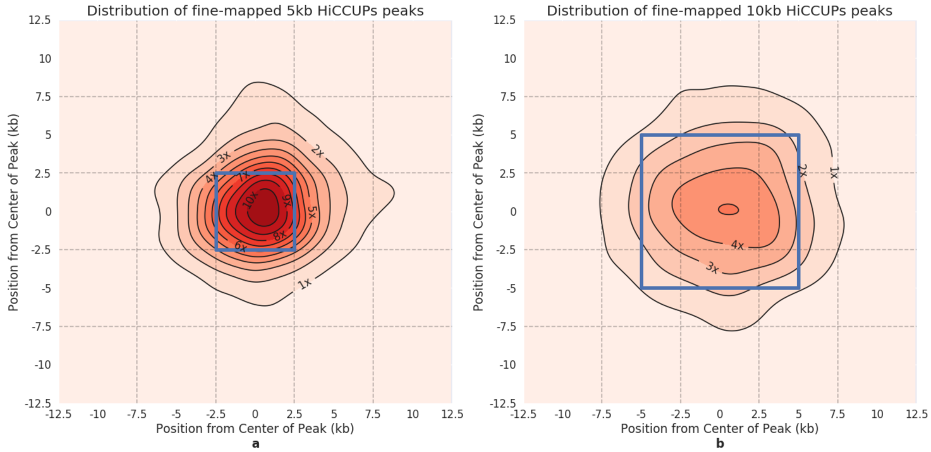

**Supplementary Figure 2. Distribution of fine-mapping predictions for different size HiCCUPs peaks.** Kernel Density Estimation (KDE) plots showing the distribution of $\chi$-SCNN's fine-mapping predictions within GM12878 peaks after extending the original peak equally in both directions to form a 25kb peak. To generate plots, we used the 'jointplot' function with the KDE option in Python's Seaborn package. **(a)** For 5kb interaction peaks extended to 25kb, fine-mapped positions are strongly concentrated around the original 5kb peak (center blue bin). Enrichment in center 5kb bin is 8.9 fold compared to random guessing. **(b)** For 10kb peaks extended to 25kb, fine-mapped positions are concentrated in the original 10kb peak (center blue bin). Enrichment in center 5kb bin is 3.5 fold. There were no peaks called at 25kb for GM12878. The positive direction on the axes points toward the exterior of the interactions. The mode of the 5kb peak plot is shifted toward the positive direction, meaning that fine-mapped peaks are most likely to be approximately 1kb further out than the center of the called originally called peak.

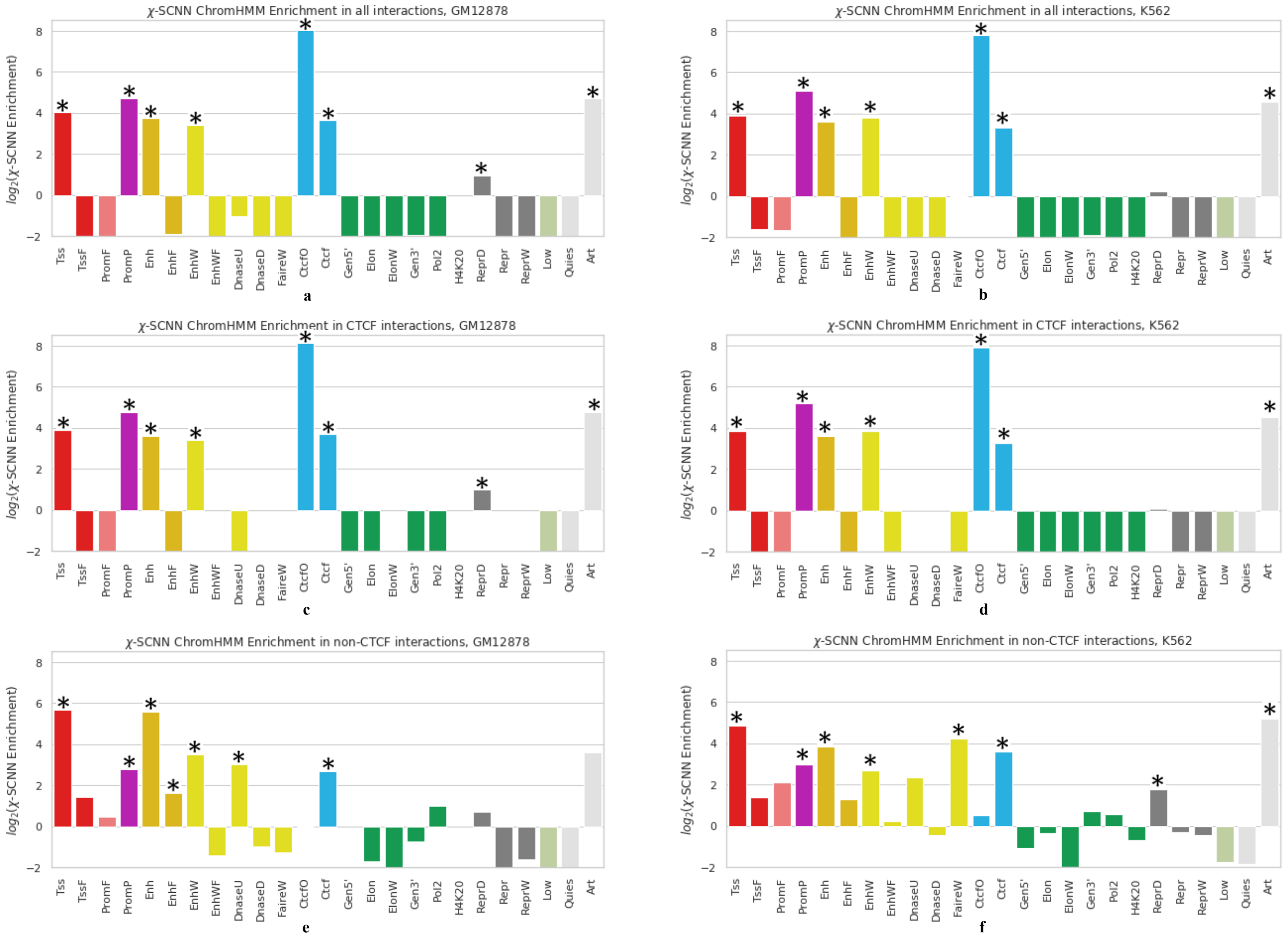

**Supplementary Figure 3. Enrichment of ChromHMM states of regions predicted by $\chi$-SCNN.** $\log_2$ fold enrichments of ChromHMM state for $\chi$-SCNN fine-mapped positions in interactions for GM12878 and K562 across (**a**, **b**) all interactions, (**c**, **d**) CTCF-associated interactions, and (**e**, **f**) non-CTCF-associated interactions. Panel (a) is the same as **Fig. 5**, but shown again for direct comparison with the other panels. Any $\log_2$ fold depletions less than -2 (1/4x) were truncated. Significant enrichments (adjusted p-value < 0.001, binomial test) are marked by an asterisk.

| Metric | | GM12878 | K562 |
|---|---|---|---|
| Maximum Matrix Resolution (kb) | | 1 | 5 |
| Peak Resolution (kb) | | 5, 10 | 5, 10, 25 |
| Training Resolution (kb) | | 25 | 25 |
| Number of Peaks, 5kb | | 6316 | 1547 |
| Number of Peaks, 10kb | | 3132 | 2343 |
| Number of Peaks, 25kb | | - | 2167 |
| Number of Peaks, total | | 9448 | 6057 |
| Number of DNase and ChIP-seq Tracks | | 100 | 148 |
| **Hyperparameter Name** | **Searched Values** | **Chosen Hyperparameter** | |
| # Encoder Kernels | $K_{Enc}$ = 12:2:32 | 26 | 16 |
| # Convolution Kernels | $K_{Conv}$ = 12:2:32 | 16 | 28 |
| Convolutional Width | $C$ = 6:2:12 | 8 | 10 |
| # Dense Kernels | $K_{Dense}$ = 12:2:32 | 16 | 16 |
| Regularization Type | $L$ = {L1, L2, L1+L2} | L1+L2 | L1+L2 |
| Regularization Strength | $S = \{0, 10^{-6}, 10^{-5}, 10^{-4}\}$ | 0 | $10^{-6}$ |
| Dropout Magnitude | $M$ = {0, 0.1, 0.25, 0.5} | 0.25 | 0 |
| **χ-SCNN Classification Performance** | | | |
| AUROC (validation) | | 0.973 | 0.974 |
| AUROC (test) | | 0.959 | 0.977 |
| AUPRC (validation) | | 0.977 | 0.974 |
| AUPRC (test) | | 0.963 | 0.972 |

**Supplementary Table 1. Detailed information on input data, hyperparameter search, and classification performance.** (**top**) For each cell type, the number of peaks separated by size and the number of available features. (**middle**) The values of the hyperparameters considered and the values chosen for each cell type. We performed a random hyperparameter search and chose the combination of hyperparameter values that yielded the highest validation AUROC. (**bottom**) The validation and test AUROCs and AUPRCs for classification. Test performance was reported on a chromosome withheld from the training and hyperparameter optimization

| 5kb Peaks | GM12878 | | | K562 | | |
|---|---|---|---|---|---|---|
| | #/6316 | % | fold | #/1547 | % | fold |
| $\chi$-SCNN (all marks) | **2260** | **36%** | **8.9** | **507** | **33%** | **8.2** |
| $\chi$-SCNN (primary marks only) | 688 | 11% | 2.6 | 148 | 10% | 2.4 |
| $\chi$-SCNN (primary+secondary marks) | 1913 | 30% | 7.6 | 392 | 25% | 6.3 |
| $\chi$-SCNN (primary+secondary marks+CTCF) | 2129 | 34% | 8.4 | 459 | 30% | 7.4 |
| CTCF | 2073 | 33% | 8.2 | 429 | 28% | 7.0 |
| RAD21 | 2165 | 34% | 8.6 | 472 | 31% | 7.6 |
| SMC3 | 2183 | 35% | 8.6 | 484 | 31% | 7.8 |
| Mean of all Signals | 1631 | 26% | 6.5 | 281 | 18% | 4.5 |
| Single Best Signal | 2183 | 35% | 8.6 | 484 | 31% | 7.8 |
| Logistic Regression on all data | 2091 | 33% | 8.3 | 442 | 29% | 7.1 |
| **10kb Peaks** | #/3132 | % | fold | #/2343 | % | fold |
| $\chi$-SCNN | **1771** | **57%** | **3.5** | **1205** | **51%** | **3.2** |
| $\chi$-SCNN (primary marks only) | 856 | 27% | 1.7 | 536 | 23% | 1.4 |
| $\chi$-SCNN (primary+secondary marks) | 1615 | 5.2% | 3.2 | 1055 | 45% | 2.8 |
| $\chi$-SCNN (primary+secondary marks+CTCF) | 1720 | 55% | 3.4 | 1172 | 50% | 3.1 |
| CTCF | 1642 | 52% | 3.3 | 1081 | 46% | 2.8 |
| RAD21 | 1673 | 53% | 3.3 | 1149 | 49% | 3.1 |
| SMC3 | 1710 | 55% | 3.4 | 1131 | 48% | 3.0 |
| Mean of all Signals | 1472 | 47% | 2.9 | 812 | 35% | 2.2 |
| Single Best Signal | 1710 | 55% | 3.4 | 1137 | 49% | 3.0 |
| Logistic Regression on all data | 1666 | 53% | 3.2 | 1109 | 47% | 3.0 |

**Supplementary Table 2. Peak recovery metrics for different size peaks, cell types, and methods.** For each cell type, the three columns represent the number of correctly recovered peaks, the percentage of recovered peaks, and the fold enrichment over expected by chance. Enrichments for 10kb peaks are lower because the expected number by chance is larger; however, the percentage correctly recovered is higher. Across all methods, $\chi$-SCNN on all marks performs the best (in bold). $\chi$-SCNN applied to primary+secondary marks and CTCF consistently performs significantly better than CTCF alone ($p$-value $< 0.05$, two proportions z-test).
